# Attention seeking in a spatially explicit game of mate choice and the evolution of dimorphic ornaments

**DOI:** 10.1101/257329

**Authors:** Szabolcs Számadó

## Abstract

The evolution of conspicuous signals fascinated biologist ever since Darwin. The Handicap Principle was dominant explanation in the last decades; it proposed that exaggerated and conspicuous signals are costly signals of quality. There are other less popular explanations however, one them is that conspicuous signals function to call the attention of potential receivers. These ‘attention seeking displays’ need not reveal the quality of the signaller. There are many empirical examples and recently the idea was modelled in terms of a simple action-response game. However, action-response games model an interaction of a pair of signaller and receiver, thus they omit potential competition between signallers, which could be a crucial force behind the evolution of attention-seeking displays. Here I model this competition in a spatially explicit model of mate choice where males can give a continuous signal to call the attention of potential mates. The results show that attention-seeking displays readily evolve to the allowed maximum when the cost of signalling is low. However, dimorphism evolves when the cost of signalling is high. The population consist of two types of males at this dimorphic state: males that do not give a signals and males that give the highest intensity signal possible. The results show that variation in quality is not a necessary requirement for the evolution of dimorphic traits.

## 1 Introduction

Conspicuous signals abound in nature. The currently dominant explanation is that these signals are ‘handicaps’: costly signals of quality [1, 2]. An alternative explanation is that these signals help receivers to find signallers. This idea was first proposed in the context of birdsong; Richards [3] found that bird song can be partitioned into an alerting and a message component in rufus-sided towhees (*Pipilo erythropthalmus*). Such signals have been found in many taxa, examples include: push up displays of *Anolis* lizard (*Anolis gundlachi*, [4]), tail-flick of Jacky dragons (*Amphibolurus muricatus*, [5]); white bones in the nest of spotted bowerbird (*Chlamydera maculata*) [6]; mating displays of satin bowerbird (*Ptilonochyncus violaceus*) [7], sage grouse (*Centrocercus urophasianus*) [8] or ring-necked pheasant (*Phasianus colchicus*) [9].

Despite the abundance of empirical examples, the modelling of the idea drew little attention in the past. Recently, Számadó [10] investigated such alert signals, which he named ‘attention-seeking-displays’ (ASD) in a simple action-response game. He was able to show that giving and listening to such displays can be evolutionarily stable. Számadó’s model [10], however, investigated the interaction of a pair of individuals, thus it omitted the competitive context of such signals. Here I incorporate competition between signallers in a spatially explicit game of attention seeking and propose a simpler model to explain the evolution of ASDs.

## 2. The model

Here I model the mate search of a sexually reproducing species. Females can mate with only one male, whereas males can mate with as many females as they can. Females have to search for partners, this search can be costly (*c*_*S*_). Males can give a signal called ‘attention-seeking-display’ (ASD) to help this search. Let’s assume that the probability of finding a partner (*p*) is proportional to the intensity of the signal (*x*). Let’s further assume that giving a signal is costly and the cost is proportional to the intensity of the signal (*c*(*x*)). There is no difference between males and females and males receive the same benefit (*b*) from mating. Then the fitness of females (*v*) and males (*u*) can be written up as follows:

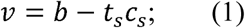

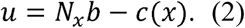

Where *t*_*s*_ denotes the time spent with mate search by the female and *N*_*x*_ denote the average number of matings a male achieves as a function of signal intensity.

I used individual based computer simulations to assess the evolutionary trajectories of different populations. The computer simulations model a population of individuals on a grid (*n*=100x100). Each grid point was occupied by a male and a female. Signaller and receiver behaviour coded by “genes” that can be inherited into the next generation.

One step of the simulation consists of two stages: (i) mate search and mating and (i) grid update. The receiver (female) behaviour was fixed in the simplest version: females searched for males at the grid position they occupied and in the direct neighbourhood of their position (8 nearest patch: Moore neighbourhood). Male strategy was coded as the intensity of the signal (*x*). Females selected from males proportional to the power of their signal intensity (*x*^*2*^). The grid was updated after all females found a male to mate with. Dispersion (0<*d*<1) was allowed at this stage, where dispersion implemented as swapping a male at a random position with a random neighbour with a probability *d*. A grid update consists of *n* steps of using the pairwise comparison rule (*PC*) [11-15]. The fitness a randomly selected individual (*i*) compared with one its neighbours (*j*). The neighbour can occupy the position of the resident with a probability *ƒ* if the fitness of the neighbour is larger than that of the resident. This probability was calculated using the Fermi function [11]:

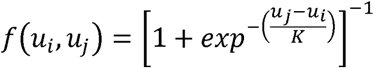

Where the constant *K* gives the steepness of the functions (*K*=1 was used in the simulations). Females competed with females and males with males to keep a constant 1:1 sex ratio. The offspring inherits its genes from its parent with a mutation probability =0.05. Simulations were iterated for *s*=1000 steps.

## 3. Results

Attention seeking displays evolve to the allowed highest intensity when the cost was low. Figure 1 shows the timeline of all runs without dispersal. There is a clear phase transition with increasing cost: the initial increase of average level of ASD shows a clear drop when *c*=18 or higher. This overshot is smaller with larger cost. The populations evolve towards a dimorphic equilibrium after this threshold (*c*=18). Individuals with 0 intensity signal start spreading in these population after the initial increase of ASD levels. At the end the population becomes dimorphic where individuals either give the maximal ASD (or close to it) or give none at all. The frequency of signallers not giving signal is increasing with increasing cost; the average level of ASD is decreasing as a result. Figures 2,3,4 and 5 show the final grid for *c*=8, *c*=18, *c*=28 and *c*=38 respectively. SI files (c_0_1400_1599.mp4, c_10_1400_1599.mp4, c_20_1400_1599.mp4, c_30_1400_1599.mp4) show the time progression of these runs on the grid. Introducing dispersal did not change the above pattern. However, the dimorphic final state evolves faster in each parameter combination with increasing dispersal. Figure 6 shows the timelines of all runs with *d*=0.25.

**Figure 1.**
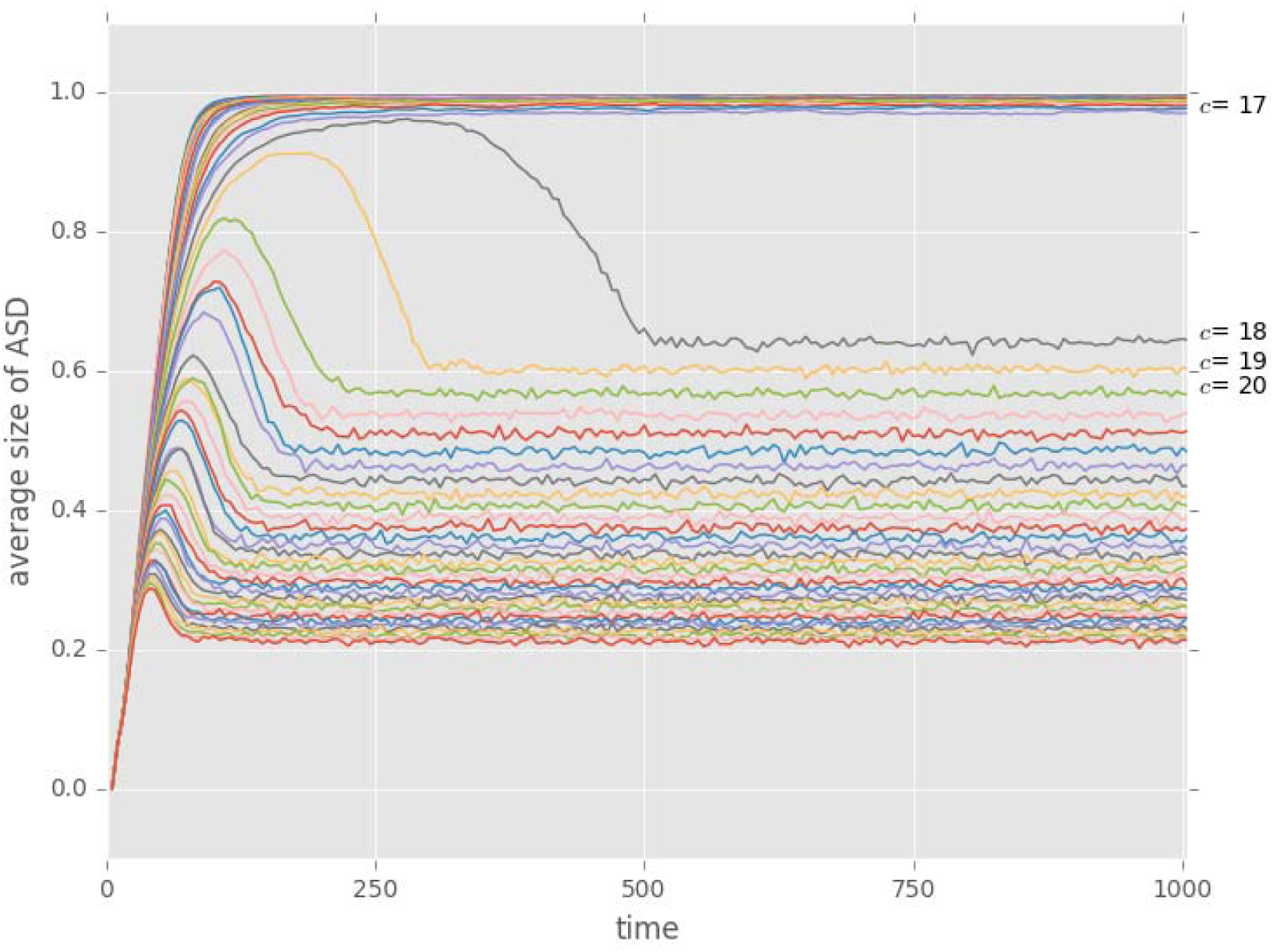
The average levels of attention seeking as a function of time for all runs without dispersal.

**Figure 2.**
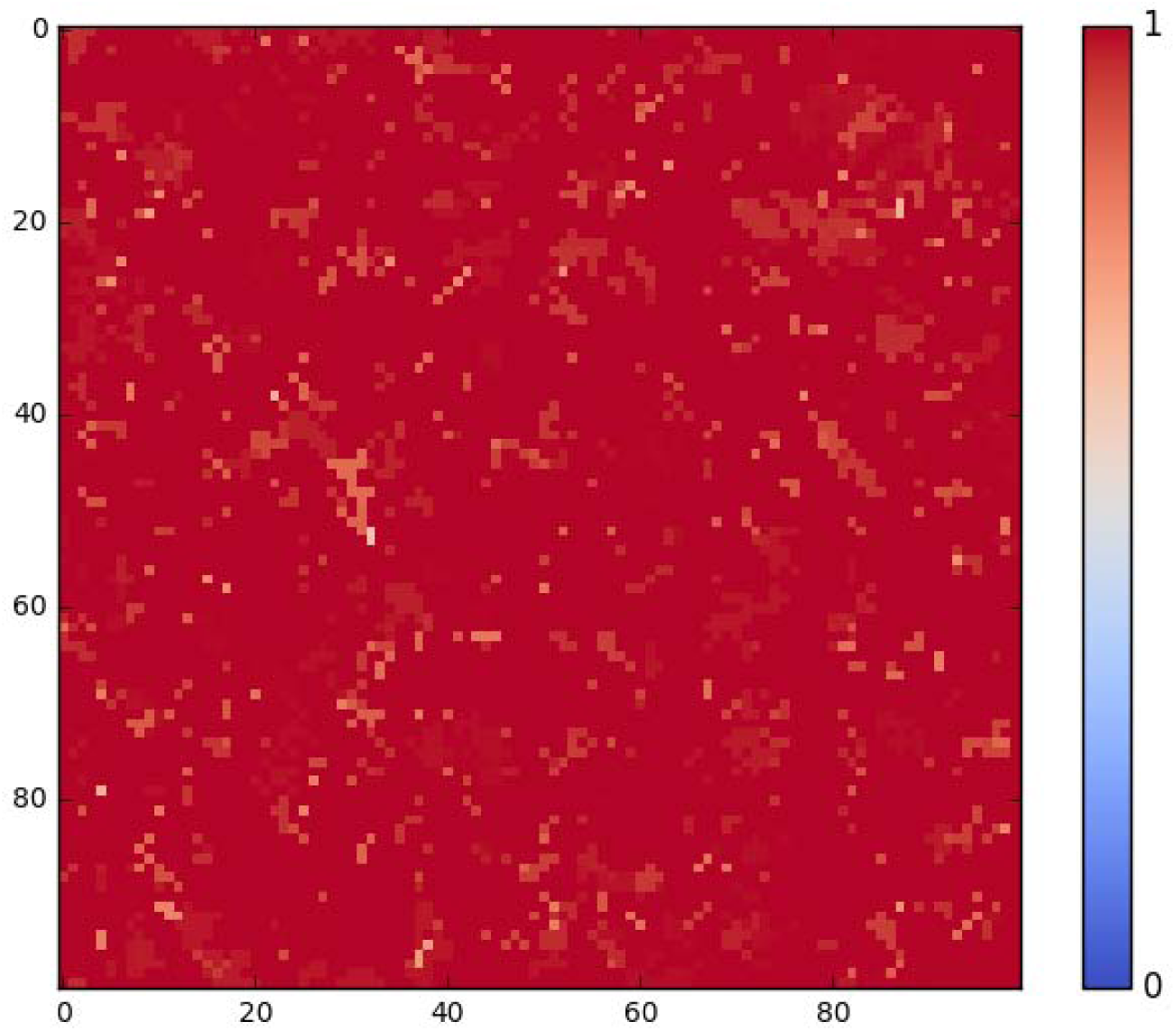
Final grid (t=1000) with *c*=8. ASD intensity increases from blue to red.

**Figure 3.**
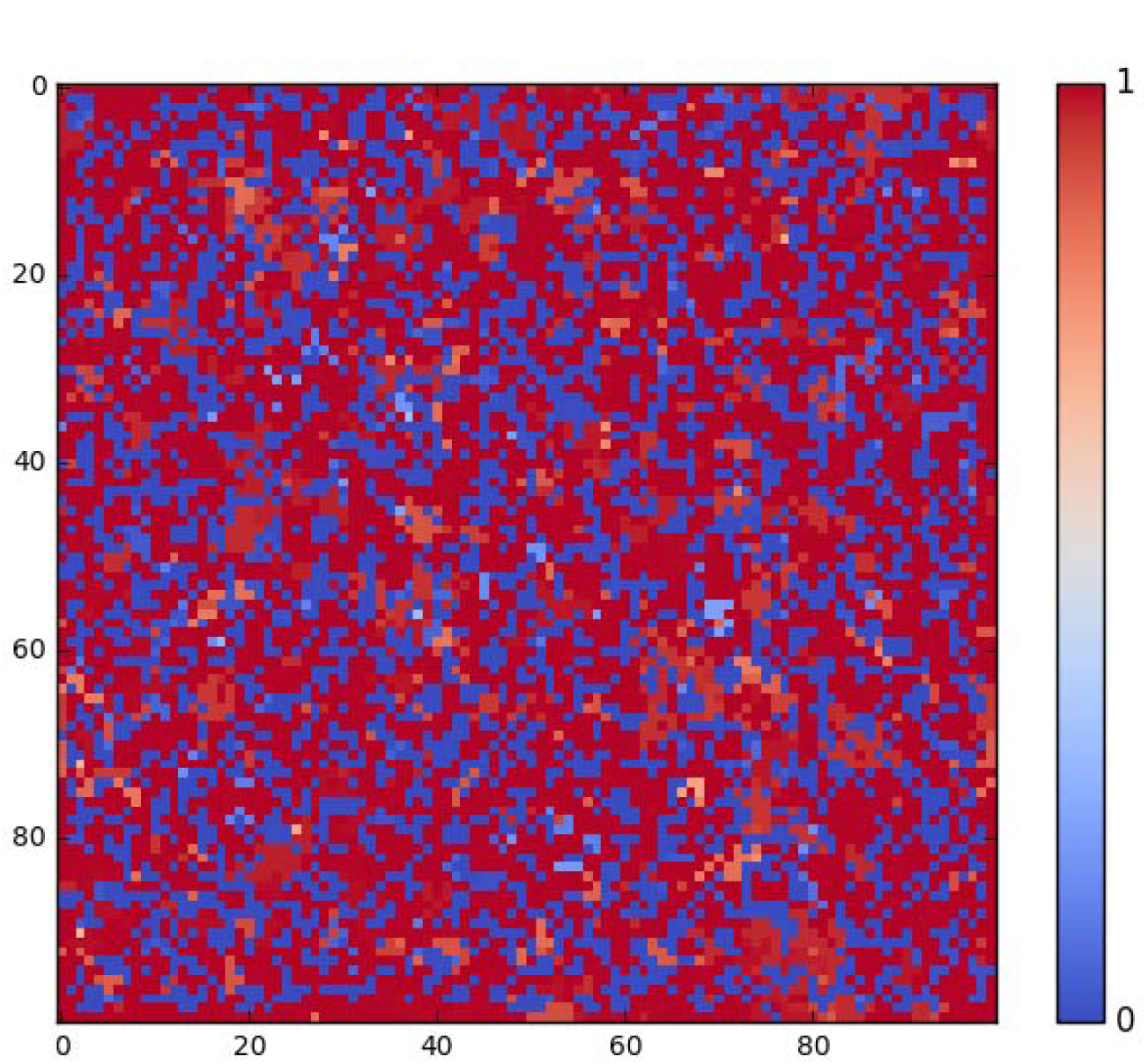
Final grid (t=1000) with *c*=18. ASD intensity increases from blue to red.

**Figure 4.**
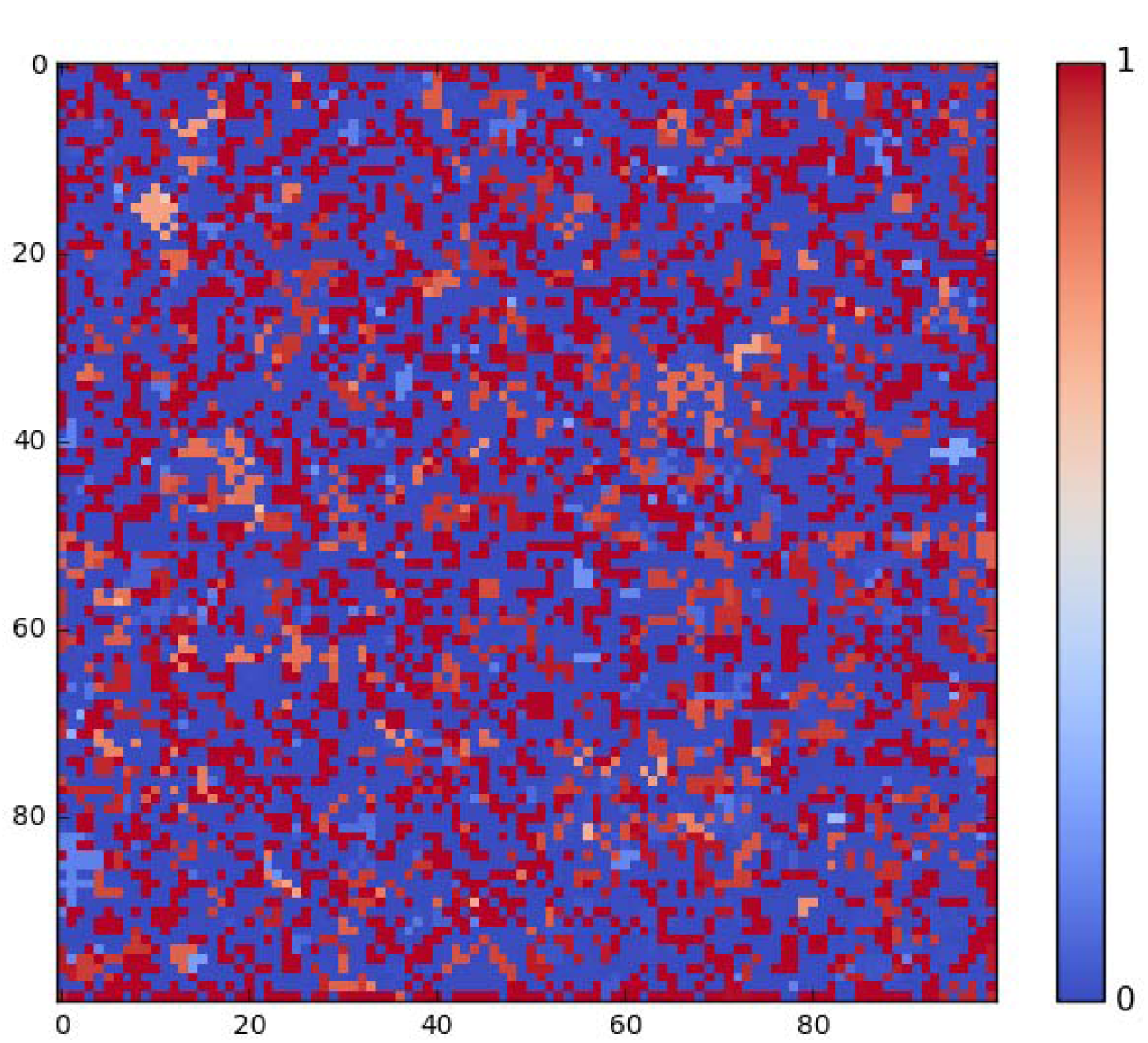
Final grid (t=1000) with *c*=28. ASD intensity increases from blue to red.

**Figure 5.**
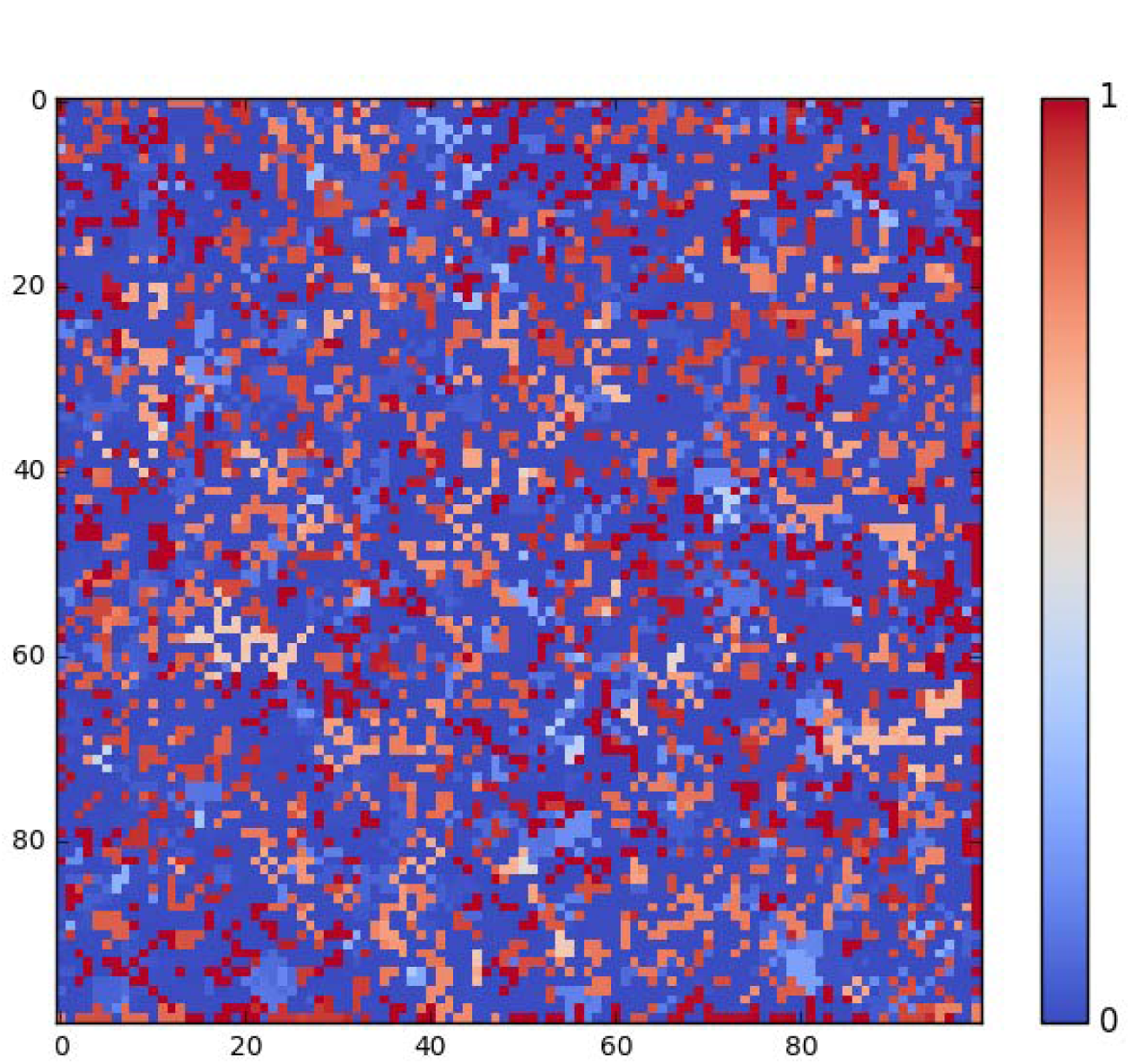
Final grid (t=1000) with *c*=38. ASD intensity increases from blue to red.

**Figure 6.**
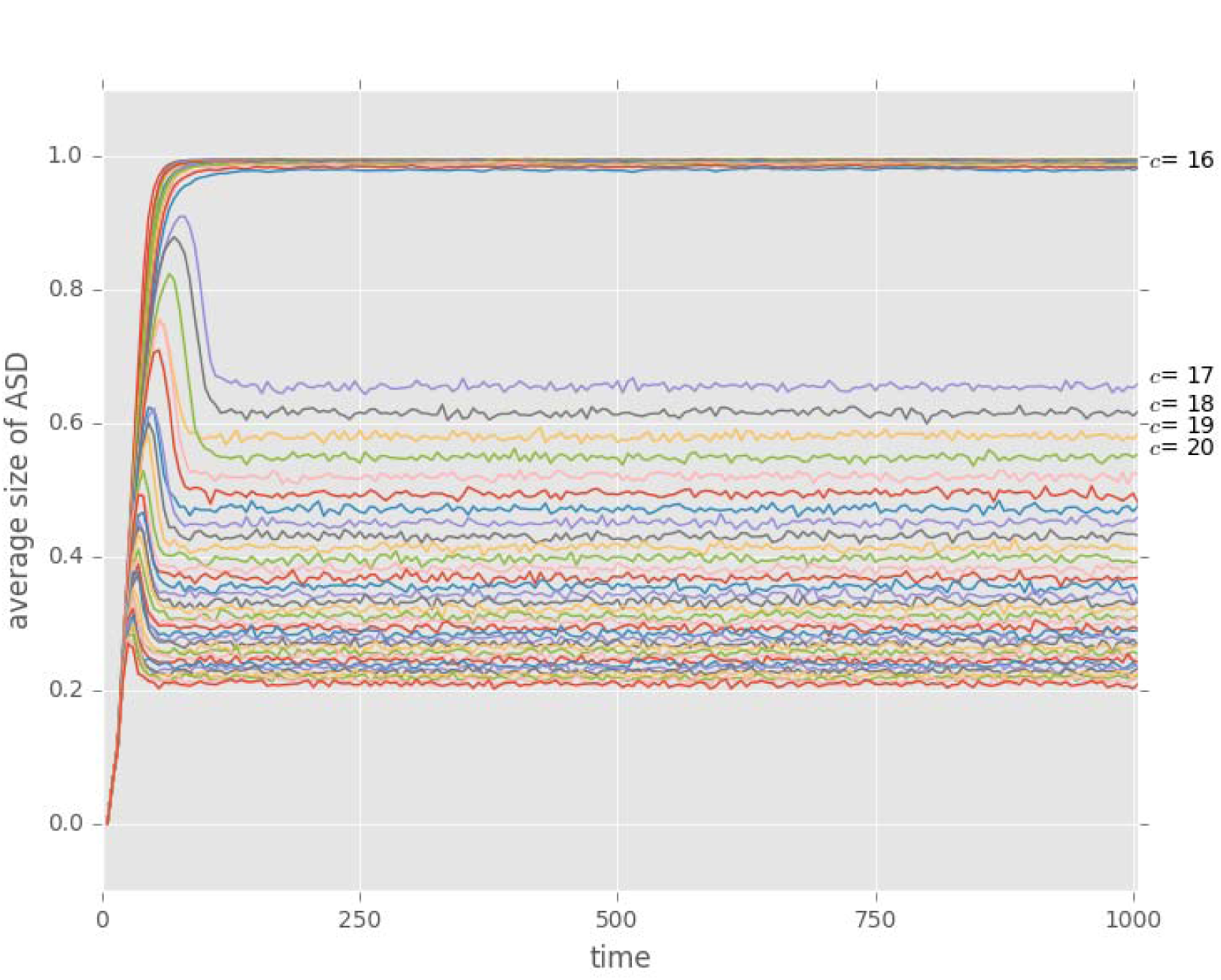
The average levels of attention seeking as a function of time for all runs with *d*=0.25.

## 4. Discussion

Attention seeking displays readily evolve when the cost of signalling is low. A dimorphic equilibrium evolves after a threshold with increasing cost. This dimorphic state evolves without quality signalling, all the males have the same quality in the model and there is no differential cost to signalling. Consequently, these signals are not ‘handicaps’ or ‘costly signals’. The results show that the key ingredient for the evolution of dimorphism is the ‘playing-the-field’ assumption, i.e. that the fitness of the individuals depends on the strategy played by others and it is not an absolute measurement.

The results strongly suggest that not all extravagant signals are handicaps. Conspicuous signal can evolve for attracting mates without revealing the quality of the signaller.

## Author contribution

S.S. conceived the idea, analysed the model and wrote the paper.

## Founding

Sz.Sz. was supported by National Research, Development and Innovation Office (NKFIH) OTKA grant K 108974 and by the European Research Council (ERC) under the European Union’s Horizon 2020 research and innovation programme (grant agreement number 648693).

